# Single snapshot quantitative phase imaging with polarization differential interference contrast

**DOI:** 10.1101/2021.06.06.447109

**Authors:** Mark Strassberg, Yana Shevtsova, Domenick Kamel, Kai Wagoner-oshima, Hualin Zhong, Min Xu

**Affiliations:** Dept. of Physics and Astronomy, Hunter College and the Graduate Center, The City University of New York, 695 Park Ave, New York, NY 10065; Biology Dept., Hunter College and the Graduate Center, The City University of New York, 695 Park Ave, New York, NY 10065

## Abstract

We present quantitative phase imaging with polarization differential interference contrast (PDIC) realized on a slightly modified differential interference contrast (DIC) microscope. By recording the Stokes vector rather than the intensity of the differential interference pattern with a polarization camera, PDIC enables single snapshot quantitative phase imaging with high spatial resolution in real-time at speed limited by the camera frame rate alone. The approach applies to either absorptive or transparent samples and can integrate simply with fluorescence imaging for co-registered simultaneous measurements. Furthermore, an algorithm with total variation regularization is introduced to solve the quantitative phase map from partial derivatives. After quantifying the accuracy of PDIC phase imaging with numerical simulations and phantom measurements, we demonstrate the biomedical applications by imaging the quantitative phase of both stained and unstained histological tissue sections and visualizing the fission yeast *Schizosaccharomyces pombe*’s cytokinesis.

---

Quantitative phase imaging (QPI) has recently emerged as a powerful tool for measuring biomass distribution and time-lapse imaging of cellular dynamics with recent advances in the spatial resolution [1], the imaging speed [2, 3], high content imaging [4], and 3D imaging [5, 6]. Typical QPI, however, has a much poorer axial resolution (~560 nm) than the lateral resoluti~350 nm) [6]. The differential interference contrast (DIC) microscope, which images the interference fringes and is only qualitative, on the contrast, can achieve excellent lateral resolution and axial discrimination. Aided by computation, quantitative DIC has emerged as one attractive quantitative phase imaging modality [7, 8], particularly the seminal work by Shribak et al. [9, 10]. Its notable advantages include no need for phase unwrapping, tolerance to scattering [11], and the optical sectioning much sharper than other QPI methods [9].

In a conventional Nomarski DIC microscope, the incident wavefront and its replica with orthogonal polarization directions pass through the same sample with an offset determined by the shear *s*, recombine and interfere. The sample phase profile is qualified by the resulting interference pattern typically after filtered by a polarization analyzer to enhance the contrast. In contrast, the proposed PDIC microscope (see Fig. 1) records the interference Stokes vector without filtering (i.e., the DIC analyzer is disengaged). In a well-aligned DIC microscope, the incident wavefront ***E***_**1**_ and its replica ***E***_2_ have equal amplitudes |***E***_1_| = |***E***_2_| = *E*_0_ and their polarization directions are ±*π*/4,respectively, away from the DIC polarizer orientation (*y*-axis). Both fields pass through the same specimen at an offset (DIC shear) *s*. After passing through the sample and combining by a second Wollaston prism, *E*_2_ accumulates a phase difference Δ = ***s*** · ∇*ϕ* + *ϕ_b_* over *E*_1_ due to both the specimen of a phase profile *ϕ* and the DIC bias *ϕ_b_*. The interference of these two fields, however, is not directly imaged inside a typical microscope. A mirror further reflects their interference ***E***_1_ + ***E***_2_ exp(*i*Δ) in our microscope to a camera attached on a side port. The *p*-polarization component reflected by the mirror experiences an extra phase delay *δ* and an unequal reflectivity (the reflectivity ratio is *r*) than the *s*-polarization component.

**Fig. 1.**
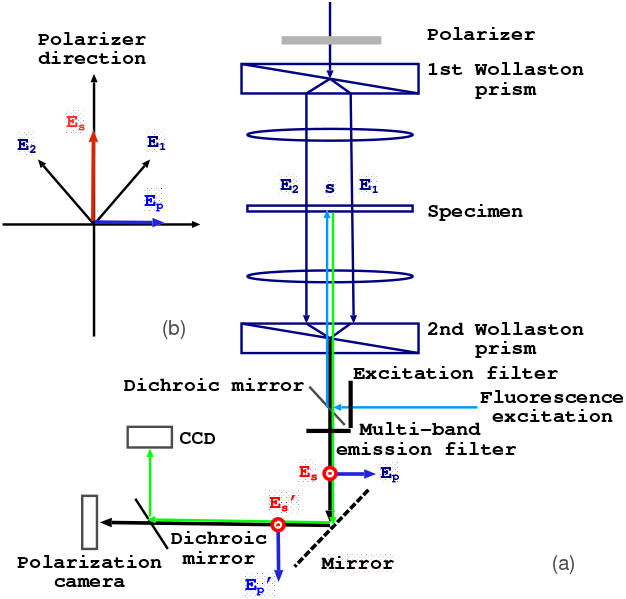
(a) A schematic diagram and (b) orientations of the directions for an inverted PDIC microscope. The first Wollaston prism splits the incident beam linearly polarized along the polarizer direction (the *y*-axis) into in-phase ***E***_1_ and ***E***_2_ polarized in the directions, 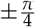 away from the *y*-axis, respectively. The DIC shear ****s**** is along the *E*_1_ direction. ***E**_s_* and ***E**_p_* are the *s*- and *p*-polarization components of the electric field reflected by the mirror. The emerging wavefronts 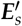 and 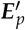 interfere and form an image at the side port recorded by a polarization camera. The epi-fluorescence optical path for simultaneous measurement is also shown.

Inside the reference frame (viewed facing the impugning beam) of the polarization camera, the net electric field is expressed as

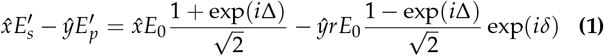

where 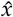 and *ŷ* are the unit vectors along the *x* and *y* directions. The unfiltered interference pattern produced by Eq. (1) is recorded by a polarization camera, yielding *I* = (*I*_0°_, *I*_45°_, *I*_90°_, *I*_135°_)^*T*^ for the intensity of light linearly polarized along 0°,45°,90°,and 135° directions, respectively, and a spatially resolved Stokes vector for the specimen given by

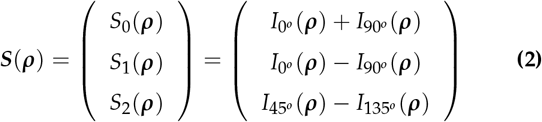

in a single snapshot where ***ρ*** **=**(*x*, *y*) are the pixel coordinates.

This provides a straightforward expression for Δ = atan2(sin Δ, cos Δ) from

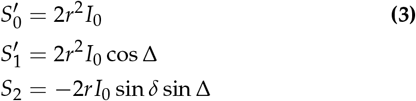

where *I*_0_ ≡ |*E*_0_|^2^ and 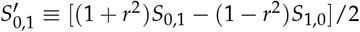 from Eq. (1). The results are particularly simple in the limit of *r* = 1. In this case, ****S**** reduces to 2*I*_0_(1, cos Δ, −sin Δ sin *δ*)^*T*^ and the variance of Δ is given by

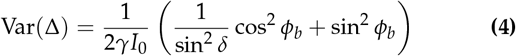

where the measured light intensities *I_i_* (*i* = 0°,45°,90°, and 135°) are assumed to be dominated by the shot noise and specified by their variances equaling to Var(*I_i_*) = *I_i_* /*γ*. The least error Var(Δ) is reached at *ϕ_b_* = *π*/2.

To shed more light to the above result, the image formation is also analyzed with the weak object transfer function (WOTF) for a telecentric microscopic imaging system under Köhler illumination [12]. Consider an in-focus thin sample whose complex transmission profile *a*(****ρ****) exp [*iϕ*(***ρ***)] = *a*_0_ + Δ*a*(****ρ****) + *ia*_0_ *ϕ*(****ρ****) where Δ*a* and *ϕ* are the amplitude and phase variations (the phase delay from the background has been assumed to be zero without losing generality). It is straightforward to show

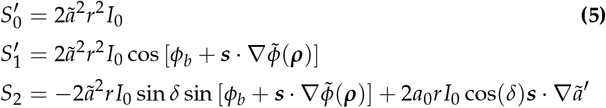

to the first order where 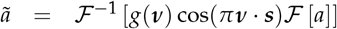, 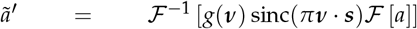, and 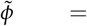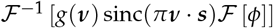 are the filtered amplitude and phase, 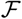 and 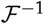 denote Fourier transform and inverse Fourier transform, sinc(*x*) ≡ sin(*x*)/*x* is the sinc function, ***ν*** is the spatial frequency, and *g*(****ν****) ≡ *C*_BF_(****ν****, 0)/*C*_BF_(0, 0) is the normalized partially coherent transfer function for a brightfield microscope [13]. According to Eq. (5), the setting of *ϕ_b_* = *π*/2 is optimal as *S′*_1_ gains the highest sensitivity 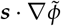 to and the influence of the unwanted second term in *S*_2_ due to absorption is minimized. The diffraction theory result (5) agrees with (3) from ray optics provided that the phase in (3) should be understood as the filtered 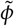 and its validity requires smooth variations in absorption 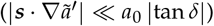. Further *I*_0_ in (3) corresponds to the transmitted light intensity 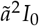 in (5).

The filtered phase map 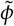 can then be obtained from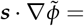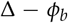. Fourier transform has been used before to find 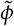 from its partial derivative [14]. An iterative approach based on total variance (TV) regularization is presented below to improve the above solution, solving the phase profile *u*(*x*, *y*) from *∂_x_u ≡ ∂u*/*∂x* = *ϕ_x_* (*x*, *y*) given the measured phase gradient *ϕ_x_* assuming ****s**** is along the *x*-axis. Specifically, we formulate it as the following optimization problem:

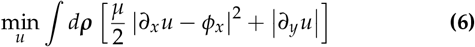

where *µ* is a positive constant and controls the degree of TV regularization. Eq. (6) can be solved using an augmented Lagrangian function [15, 16]

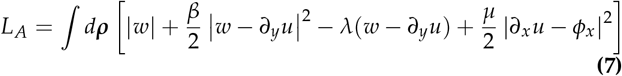

with the alternative direction method (ADM),

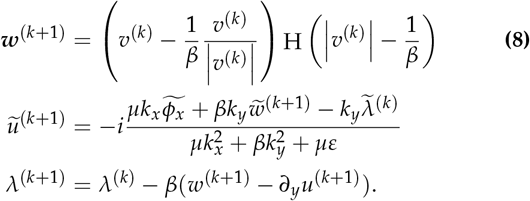

Here 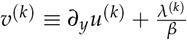, H is the Heaviside function, the superscript (*k*) means the quantities at the *k*-th iteration, 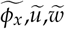, and 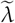 are the Fourier transforms of *ϕ_x_*, *u*, *w*, and *λ*, respectively, *k_x_* and *k_y_* are the *x*- and *y*-components of the wavenumber inside the Fourier space, and ɛ = 10^−6^ is introduced to avoid the division by zero. We initiate the iterative solution by setting

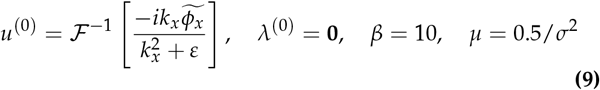

where **0** is a zero matrix of the same shape as *ϕ_x_*, and 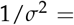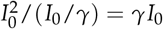 is a parameter representing the signal to noise ratio of the measurement. The initial solution *u*^(0)^ is provided by the Fourier transform solution to the partial derivative [14]. The final solution is reached when the relative change in *u* is less than a threshold (10^−3^) or the image quality (sharpness) starts to drop. The genuine phase map *ϕ* is obtained from 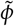 by deconvolution. The halo effect is removed computationally [17] when needed. The stability of the recovered phase is approximately the same as the variance of *ϕ_x_*, i.e., Var(*u*) ≃ Var(*ϕ_x_*) ≃ (2*γI*_0_)^−1^.

We then perform simulations and experiments to quantify the accuracy of PDIC phase imaging and demonstrate the biomedical applications. Experiments were performed on an inverted epifluorescence DIC microscope (IX73, Olympus). The light source is a Halogen 100W lamp filtered by a Green filter (central wavelength 0.545*µm*). The numerical apertures for the condenser and the water immersion UPLANSAPO 60 objective are 0.55 and 1.2, respectively. The microscope is equipped with a focus drive and motorized scan system (ASI) and a dual-camera adaptor (TwinCam, Cairn). The PDIC and fluorescence images are recorded, respectively, by a polarization camera (Blackfly S Polarized, FLIR) and a digital color camera (Blackfly S Color, FLIR). PDIC and fluorescence images are co-registered and can be captured simultaneously. The exposure time is set at 10 ms. The system parameters have been measured separately with *r* = 0.981, *δ* = −63.2°, *s* = 0.22*µm*, and the pixel size 0.0564*µm* from a set of calibration experiments. To correct the potential phase artifacts introduced by optical elements inside the microscope, we measure Δ_bg_ for a uniform background under the identical condition and obtain 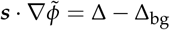. The total computation time for one phase image is ~20*s* in Matlab using one single 2.2GHz CPU.

First simulations were performed using microlith [18], which accurately predicts three-dimensional images of a thin specimen observed with any partially coherent imaging system and is modified for PDIC. Figure 2(a) shows the measured light intensities polarized along 0°,45°,90° and 135° directions for an etched SiO_2_ phase pattern of thickness 300nm imaged under a PDIC configuration (*λ* = 0.532*µm* in vacuum, numerical apertures NA_*o*_ = 1.2 and NA_*c*_ = 0.55 for a water immersion objective and a condenser, respectively, and *ϕ_b_* = *π*/2). The phase pattern contains a set of phase rectangles of width 1,0.5, 0.5, 0.25, 0.25, 0.15, and 0.15*µm*. The original phase pattern and the recovered phase patterns under low noise (0.5% noise), inhomogeneous absorption and low noise (mean amplitude reduction 25% with a standard deviation 11% and 0.5% noise), and moderate noise (3% noise) conditions are shown in Figure 2(b-e) with their horizontal profiles displayed in Figure 2(f). The recovered phase value agrees very well with the theoretical one (0.46 radian). The two smallest rectangles with a separation of 0.15*µm* are also resolved, confirming the spatial resolution limit, *λ*/(NA_*o*_ + NA_*c*_) = 0.3*µm*. The performance of the phase recovery is tolerant to both moderate inhomogeneous absorption and noise whereas the accuracy of the recovered phase deteriorates for the structures of size smaller than the spatial resolution limit.

**Fig. 2.**
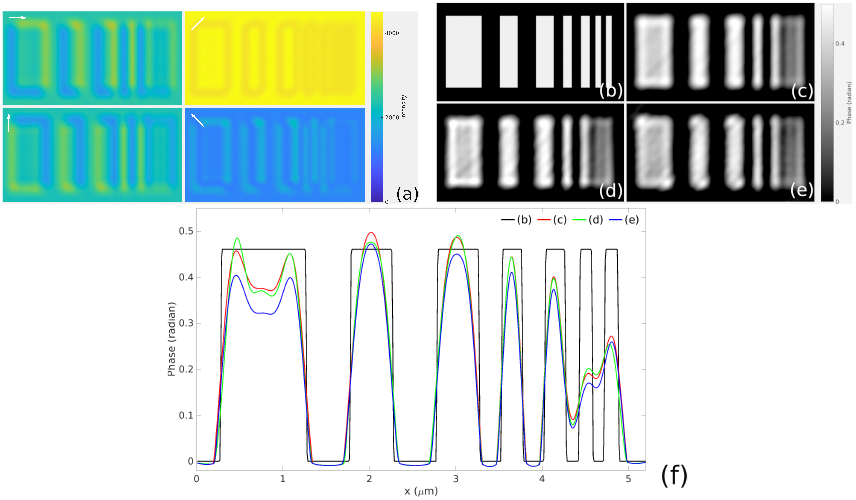
(a) The measured light intensities polarized along 0°,45°,90° and 135° directions for a SiO_2_ phase pattern of thickness 300nm. *I*_135°_ is multiplied by five for clarity. (b) The original phase pattern and the recovered phase patterns under (c) low noise (0.5% noise), (d) inhomogeneous absorption and low noise (mean amplitude reduction 25% with a standard deviation 11% and 0.5% noise), and (e) moderate noise (3% noise) conditions. (f) Their horizontal profiles.

A transmission grating (LG25-03, Thorlabs) of 300 grooves/mm and 17.5° groove angle was then imaged with the PDIC microscope under the water immersion 60× objective. Figure 3 shows the recovered phase image and physical geometry for the transmission grating. The recovered maximum phase is in excellent agreement with the nominal value (2.08 radian). The physical geometry of the grating is then deduced from the phase profile (the refractive index of the glass is *n* = 1.52) with a groove angle 19.7° ± 0.8° and a pitch of 3.30 ± 0.08*µm*.

**Fig. 3.**
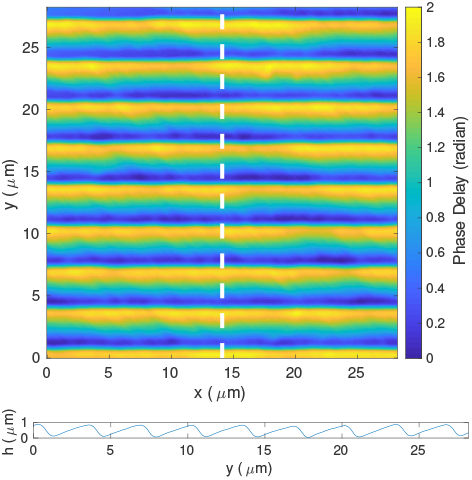
The GT25-03 transmission grating: recovered phase image (top) and physical geometry deduced from the phase profile along the dashed line (bottom).

After quantifying the accuracy of PDIC phase imaging with numerical simulations and phantom measurements, we demonstrate two biomedical applications for mapping the quantitative phase of histological tissue sections and visualizing live cells coregistered with fluorescence. Figure 4 shows the measured phase images for the hematoxylin and eosin (H&E) stained and unstained serial cuts of normal prostate tissue. The phase images for the stained and unstained serial cuts are overall similar. However, slight elevation of the phase is noticeable for the H&E stained section, especially in the areas surrounding some stromal regions.

**Fig. 4.**
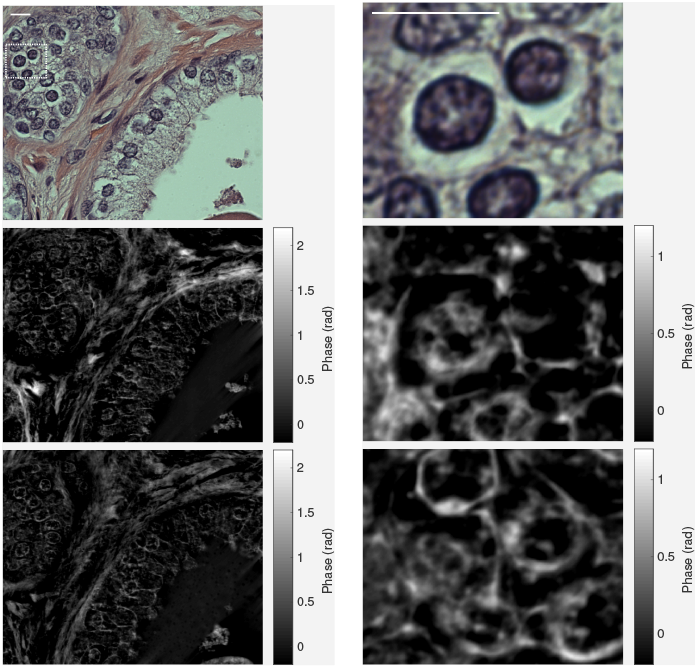
(Left) The H&E stained histologic image, the measured phase images for the H&E stained and unstained serial cuts of normal prostate tissue. (Right) The magnified images for the region highlighted by the white rectangle. Scale bar: 10*µm*.

Live cell quantitative phase imaging of the fission yeast *Schizosaccharomyces pombe* coregistered with fluorescence is shown in Figure 5. Cells were immobilized by mounting onto an agar pad containing essential nutrients. The fission yeast strain (*nup211-GFP::ura4+ura4-D18 leu1-32 ade6*) expresses the nuclear pore protein nup211 fused with the green fluorescent protein (nup211-GFP) [19], which marks the position of the nucleus. The fluorescence images were cleaned by the iterative Poisson denoising with variance-stabilizing transformations (VST) [20] and overlaid on the phase map. The cytokinesis process of cell #4 is clearly shown, including the formation and the cleavage of the septum. We also observed the fast movements of structures in the cytoplasm. Further investigations are needed to identify these subcellular structures and elucidate their functions.

**Fig. 5.**
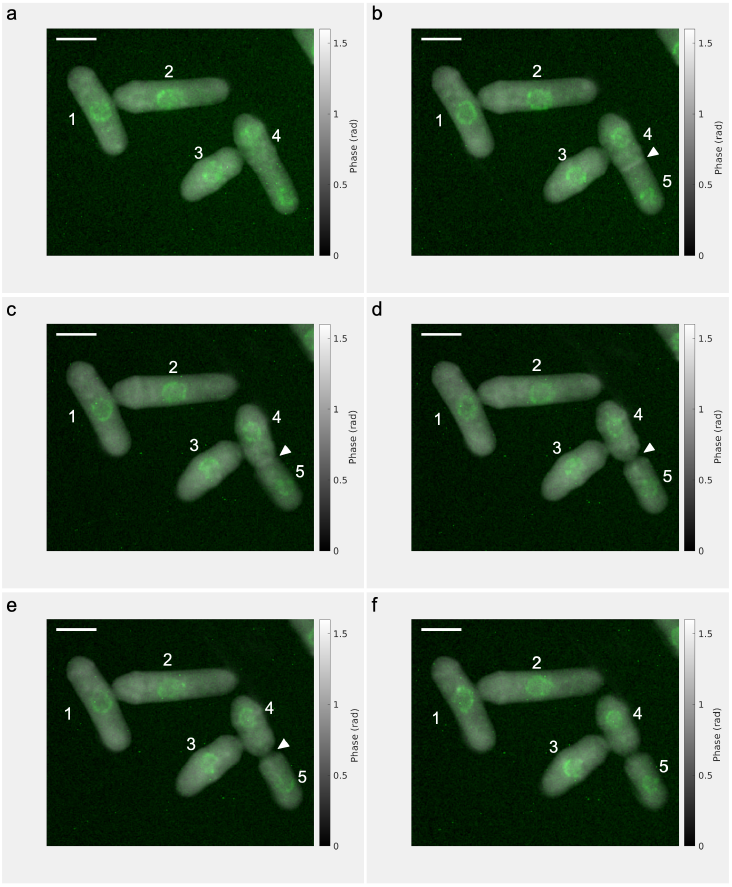
Live cell imaging of the fission yeast *Schizosaccharomyces pombe* at 3, 11, 22, 27, 30, and 34 mins in (a-f) (See Visualization I). This strain expresses the nuclear pore protein nup211 fused with the green fluorescent protein (nup211-GFP), marking the position of the nucleus. Arrowheads point to the septum, where cytokinesis occurs. Scale bar: 5*µm*.

QPI system is defined by the compromise of four figures of merit: space–bandwidth product, time-bandwidth product, spatial phase sensitivity, and temporal phase sensitivity [21]. While it is difficult to maximize each of these parameters in one single system, the PDIC microscope performs well in all four figures of merit with single snapshot broadband illumination and common-path geometry. The phase sensitivity can be estimated to be 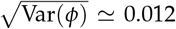 and the optical path length sensitivity 1.0nm at *λ* = 0.545*µm* from *I*_0_ ≃ 20000 (*γ* = 0.176 for the polarization camera), agreeing well with the experimentally measured spatial and temporal sensitivity of 1.15 ± 0.05nm. The phase sensitivity can be significantly enhanced by using a camera of an extremely deep well and/or by averaging over multiple images and performing spatially low pass filtering [22] when the phase error is dominated by photon noise. The phase difference measured by PDIC is inherently between two polarization states (***E***_**1**_and ***E***_2_) and hence affected by birefringence if present. Cautious interpretation is needed for strongly birefringent specimens. Compared to earlier studies [3], this work presented the PDIC quantitative phase imaging of both absorptive and transparent samples and an iterative method to improve the quality of the recovered phase map. The polarization alteration due to the mirror reflection is explicitly considered, critical for an accurate phase recovery. We derive the formalism for the PDIC microscope using both the geometrical optics and the weak object transfer function. The latter reduces to the former and reveals its validity condition that the phase is the filtered version and the variation in sample absorption is spatially smooth. Finally we note the current PDIC implementation is limited to measure the phase gradient along one single axis (the shear direction) and regularization plays an important role in phase recovery. Future studies are warranted to investigate the optimal regularization for a complex phase distribution.

In summary, we have presented quantitative phase imaging with polarization differential interference contrast (PDIC) by recording the Stokes vector rather than the intensity of the differential interference pattern with a polarization camera. PDIC enables single snapshot quantitative phase imaging of either absorptive or transparent samples with high spatial resolution in real-time at speed limited by the camera frame rate alone. The approach can integrate simply with fluorescence imaging for coregistered simultaneous measurements. We have demonstrated the accuracy of PDIC phase imaging and the biomedical applications in imaging the quantitative phase of histological tissue sections and visualizing the cytokinesis process of the fission yeast. PDIC may emerge as an attractive quantitative phase imaging method for biology and medicine.

## Supporting information

Derivation of Equations 3 and 4

## Supplemental document

See Supplement 1 for supporting content.

