## Supplementary material for "Single snapshot quantitative phase imaging with polarization differential interference contrast": Derivation of Equations 3 and 4

### 1. DERIVATION OF EQ. (3)

The recorded Stokes vector is

$$S = |E_0|^2 \begin{pmatrix} 1 + r^2 + (1 - r^2)\cos\Delta \\ 1 - r^2 + (1 + r^2)\cos\Delta \\ -2r\sin\Delta\sin\delta \end{pmatrix} \tag{S1}$$

according to Eq. (1) in the main text. The equation (3) in the main text is obtained by introducing $I_0 \equiv |E_0|^2$ and $S'_{0,1} \equiv \left[(1 + r^2)S_{0,1} - (1 - r^2)S_{1,0}\right]/2$.

### 2. DERIVATION OF EQ. (4)

The variance of $\Delta$ can be estimated using a homogeneous specimen. In this case, the measured light intensities are given by

$$\begin{aligned} I_{0^o} &= I_0(1 + \cos\phi_b) \\ I_{90^o} &= I_0 r^2(1 - \cos\phi_b) \\ I_{45^o} &= \frac{1}{2}I_0\left[1 + r^2 + (1 - r^2)\cos\phi_b - 2r\sin\phi_b\sin\delta\right] \\ I_{135^o} &= \frac{1}{2}I_0\left[1 + r^2 + (1 - r^2)\cos\phi_b + 2r\sin\phi_b\sin\delta\right] \end{aligned} \tag{S2}$$

with an error specified by their variances equaling to $\mathrm{Var}(I) = I/\gamma$.

At the limit of $r = 1$, we have

$$\begin{aligned} \cos\Delta &= \frac{1}{S_0}S_1 = \frac{2(I_{0^o} - I_{90^o})}{I_{0^o} + I_{45^o} + I_{90^o} + I_{135^o}} \\ \sin\Delta &= \frac{1}{-\sin\delta S_0}S_2 = -\frac{2}{\sin\delta}\frac{I_{45^o} - I_{135^o}}{I_{0^o} + I_{45^o} + I_{90^o} + I_{135^o}}. \end{aligned} \tag{S3}$$

We can compute

$$\begin{aligned} \mathrm{Var}(\cos\Delta) &= 4\gamma^{-1}\left(\frac{I_{0^o} - I_{90^o}}{I_{0^o} + I_{45^o} + I_{90^o} + I_{135^o}}\right)^2 \\ &\times\left[\frac{I_{0^o} + I_{90^o}}{(I_{0^o} - I_{90^o})^2} + \frac{I_{0^o} + I_{45^o} + I_{90^o} + I_{135^o}}{(I_{0^o} + I_{45^o} + I_{90^o} + I_{135^o})^2} - 2\frac{1}{I_{0^o} + I_{45^o} + I_{90^o} + I_{135^o}}\right] \\ &= \frac{1}{2\gamma I_0}(1 - \frac{1}{2}\cos^2\phi_b) \end{aligned} \tag{S4}$$

and

$$\begin{aligned} \sin^2\delta\,\mathrm{Var}(\sin\Delta) &= 4\gamma^{-1}\left(\frac{I_{45^o} - I_{135^o}}{I_{0^o} + I_{45^o} + I_{90^o} + I_{135^o}}\right)^2 \\ &\times\left[\frac{I_{45^o} + I_{135^o}}{(I_{45^o} - I_{135^o})^2} + \frac{I_{0^o} + I_{45^o} + I_{90^o} + I_{135^o}}{(I_{0^o} + I_{45^o} + I_{90^o} + I_{135^o})^2} - 2\frac{1}{I_{0^o} + I_{45^o} + I_{90^o} + I_{135^o}}\right] \\ &= \frac{1}{2\gamma I_0}(1 - \frac{1}{2}\sin^2\delta\sin^2\phi_b). \end{aligned} \tag{S5}$$

The accuracy of the recovered $\Delta$ is then $\text{atan2}(\sin\Delta, \cos\Delta) \pm \sqrt{\text{Var}(\Delta)}$ with

$$\text{Var}(\Delta) = \frac{1}{2\gamma I_0}\left(\frac{1}{\sin^2\delta}\cos^2\phi_b + \sin^2\phi_b\right) \tag{S6}$$

given in Eq. (4) of the main text.
